# Bilingual Exposure and Sex Shape Developmental Trajectories of Brain Responses to Speech-Sound Features in Infants

**DOI:** 10.1101/2024.12.02.626326

**Authors:** Marta Puertollano, Natàlia Gorina-Careta, Siham Ijjou-Kadiri, Alejandro Mondéjar-Segovia, María Dolores Gómez-Roig, Carles Escera

## Abstract

As the auditory brain becomes functional during the third trimester of pregnancy, both biological and environmental processes begin shaping its maturation, influencing how speech sounds are perceived. Biological factors, such as sex, introduce early genetic differences, while environmental experiences, like bilingualism, modulate the auditory input that infants receive. Although existing research highlights the impact of sex and bilingualism on the development of speech perception, the neural mechanisms remain unclear. In this study, we recorded frequency-following responses (FFRs) longitudinally, at birth, six months, and twelve months of age in 73 infants exposed to varying degrees of bilingual input. We modeled the developmental trajectories for neural encoding of voice pitch and speech formant structure, finding significant maturation during the first six months, followed by stabilization through the first year. Distinct developmental patterns emerged as a function of sex and bilingualism, revealing their influence on neural attunement to key speech-sound features. Bilingual exposure notably predicted lower neural pitch encoding values at six months, but higher values by twelve months. A positive effect of bilingualism on speech formant encoding was observed throughout the first year. These findings reveal how biological and environmental factors contribute to individual variability in early auditory development and speech acquisition.

**Highlights:**

- Bilingualism and sex impact early developmental trajectories in neural speech encoding
- Results suggest a female advantage in early neural encoding of speech-sound features
- Bilingual exposure positively modulates infant neural speech encoding

## 1. Introduction

Speech acquisition initiates at a very early developmental stage, as the auditory brain is already functional and able to process sounds since the beginning of the third trimester of pregnancy (Hepper & Shahidullah, 1994; Moore & Linthicum, 2007; Querleu et al., 1988; Ruben, 1995). It is around the 27th week of gestation that hearing becomes fully functional and the first fetal responses to sounds can be registered (Draganova et al., 2018; Schneider et al., 2001). At this fetal stage, the cochlea and the temporal lobe are formed and myelination appears through the brainstem and up to the auditory thalamus (Lavigne-Rebillard & Bagger-Sjöbäck, 1992; Moore et al., 1995; Moore & Linthicum, 2007). Although the exact acoustic features reaching the fetus remain unclear, intrauterine recordings suggest that the acoustic signal is altered by the maternal womb, which attenuates around 30dB frequencies above 500 Hz (Abrams et al., 2000; Gerhardt & Abrams, 1996, 2000). This filtering primarily preserves the prosodic features of speech, conveying the variations in pitch, loudness and rhythm, while suppressing the phonemic contrast information (Moon, 2017; Querleu et al., 1988).

It is during this early period that fetuses are first exposed to an acoustic environment, which significantly influences the development of their acoustic capacities (Arenillas-Alcón et al., 2023; Gorina-Careta et al., 2024; Moon et al., 2013; Partanen et al., 2013b). For instance, daily music exposure during pregnancy has been shown to positively impact the neural encoding of speech sounds at birth (Arenillas-Alcón et al., 2023). Similarly, prenatal exposure to a bilingual environment affects the neonatal acoustic sensitivity to speech frequencies (Gorina-Careta et al., 2024). Neonates also demonstrate distinct preferences that indicate prenatal acoustic learning and tuning to the prosody of their native language (DeCasper & Fifer, 1980; Fernald & Simon, 1984; Granier-Deferre et al., 2011; Moon et al., 1993, 2013). Indeed, prenatal exposure to speech evoke enduring changes in neural dynamics that further support learning and memory (Mariani et al., 2023).

Shortly after birth, infants exhibit sensitivity to a wide range of linguistically significant distinctions (Gervain & Mehler, 2010; Martinez-Alvarez et al., 2023). Newborns are able to encode the pitch of speech sounds in an adult-like manner (Arenillas-Alcón et al., 2021) and discriminate between languages they have not been exposed to, provided those languages differ rhythmically (Byers-Heinlein et al., 2010; Mehler et al., 1988; Nazzi et al., 1998). However, language acquisition relies on the capacity to classify similar yet non-identical sounds into either different or equivalent phonetic categories according to the specific language, which is dependent on further postnatal linguistic exposure (Kuhl et al., 1992, 2003; Rivera-Gaxiola et al., 2005). By the age of six months, infants typically begin to perceive the variability inherent in each phonetic unit, which enables them to identify vowels typical of their mother tongue and alters their phonetic perception toward a native-like model (Kuhl et al., 1992; Maye et al., 2002). These developmental changes are reflected in a pronounced enhancement in neural encoding of speech sound features (Ribas-Prats et al., 2023b; Puertollano et al., 2024) and coincide with their first articulation of consonant-vowel sounds, marking the babbling stage (Oller, 1992).

Exposure to distinct linguistic environments further shape developmental trajectories for infant speech processing. Monolingual infants demonstrate precise discrimination among a wide range of native and non-native phonemic contrasts by seven months of age (Best & McRoberts, 2003; Cheour et al., 1998; Rivera-Gaxiola et al., 2005). Conversely, bilingual infants at the same age do not exhibit equivalent levels of native phoneme discrimination, with within-group variability depending on the amount of exposure to each language (Best et al., 2016; Garcia-Sierra et al., 2011). As infants approach their first year of life, their speech perception undergoes an experience-driven perceptual narrowing, leading to a refined attunement to their native phoneme repertoire (Cheour et al., 1998; Kuhl et al., 2006; Werker et al., 1981) and typically aligning with their first word productions (Fenson et al., 1994). As their expertise in native phoneme contrasts develops, the ability to discriminate non-native phonemes diminishes, becoming minimal by the end of the first year (Rivera-Gaxiola et al., 2005; Tsao et al., 2006; Werker & Tees, 1984). Notably, the timing of these developmental milestones is influenced by the balance of language input, regardless of whether infants are raised in monolingual or bilingual settings (García-Sierra et al., 2011, 2016).

Alongside linguistic environments, sex is a significant biological factor influencing speech processing. Beginning in the second trimester of pregnancy, the extensive placental transmission of sex-steroid hormones shapes early speech development (Lust et al., 2010; Lutchmaya et al., 2001; Schaadt et al., 2015; Wermke et al., 2014). As early as one month after birth, female infants tend to outperform males in phonological discrimination (Friederici et al., 2008), a difference mediated by sex hormone levels that are also linked to articulatory skills at five months (i.e., babbling; Quast et al., 2016), and to later language abilities in childhood (Hollier et al., 2013; Schaadt et al., 2015). Additionally, vocabulary growth rates have been observed to be faster in female infants (Dailey & Bergelson, 2023), while male infants face a higher risk of experiencing language delays within the first three years of life (Whitehouse et al., 2012). Despite these findings, there is still limited research exploring how sex modulates the developmental trajectories of neural speech encoding during the first year of life.

Advancements in utilizing infant brain potentials have significantly pushed the study of neural mechanisms involved in speech perception and acquisition (Hervé et al., 2022). The frequency-following response (FFR) stands out as an auditory evoked potential that captures neural synchronization along the auditory pathway in response to complex sounds such as speech and music (Coffey et al., 2019; Gorina-Careta et al., 2021). FFR recordings have proven to be a powerful tool for investigating the neural encoding of speech sound features, such as pitch and fine spectrotemporal details that underlie phoneme perception (Gorina-Careta et al., 2022; Krizman & Kraus, 2019). In infancy, the FFR has been employed to characterize typical and atypical development of neural speech encoding (Banai et al., 2005, 2009; Cunningham et al., 2001; Puertollano et al., 2024; Ribas-Prats et al., 2019, 2022, 2023b, 2023a). It has also been used to study biological and environmental influences on auditory processing, including sex-related differences throughout development (Krizman et al., 2019, 2020), as well as the impact of prenatal bilingual experiences in neonates (Gorina-Careta et al., 2024), and postnatal experiences in children (Krizman et al., 2015) and adults (Skoe et al., 2017). However, these crucial factors remain unexplored across the first year of life.

The present study aimed to uncover how age, perinatal bilingual experience, and sex collectively shape neural speech-encoding mechanisms during the first year of life. To this end, we recorded FFRs from infants with varying degrees of bilingual exposure and examined their developmental trajectories regarding the neural encoding of voice pitch and formant structure content. Building on previous research depicting neural encoding achievements during the first six months of life, which stabilize by the first year (Puertollano et al., 2024), we anticipated observing similar age-related effects. Additionally, we sought to relate our findings to existing literature that highlights distinct developmental trajectories in phoneme discrimination associated with monolingual and bilingual experiences during infancy. We also expected to find sex-related differences in infant neural encoding of speech sounds, consistent with previous studies on speech processing variations.

## 2. Methods

### 2.1 Participants

73 healthy-term neonates (38 females; mean gestational age at birth = 39.73 ± 0.97 weeks; mean birth weight = 3288.8 ± 302.6 grams; mean age at evaluation = 1.62 ± 0.94 days after birth) were recruited at the Sant Joan de Déu Barcelona Children’s Hospital (Catalonia, Spain) and followed-up at six (aged 5.36 to 7.40 months after birth; mean = 6.23 ± 0.37 months) and twelve months of age (aged 11.81 to 13.22 months after birth; mean = 12.39 ± 0.36 months).

All infants participating in this study were born at term after low-risk gestations, with an adequate birth weight for their gestational age (Figueras & Gratacós, 2014). Any diagnosed pathology or an Apgar score below 7 at 1 and 5 minutes after birth were considered as exclusion criteria. None of the infants presented any risk factors for hearing impairment, as per the Joint Committee on Infant Hearing guidelines (2019). As part of the standard medical routine to ensure auditory pathway integrity at birth, all neonates had passed the universal hearing screening test based on an automated auditory brainstem response system (ALGO 3i, Natus Medical Incorporated, San Carlos, CA).

Approval from the Bioethics Committee of SJD Barcelona Children’s Hospital (Internal review board ID: PIC-185-19) was obtained for this study. Prior to the infant data collection, all parents or legal guardians signed an informed consent in accordance with the Code of Ethics of the World Medical Association (Declaration of Helsinki). All data from this study is available upon inquiry to the corresponding author.

### 2.2 Language Exposure Measurement

Infants’ prenatal linguistic exposure was evaluated through a retrospective questionnaire delivered to their mothers, as reported previously by our group (Gorina-Careta et al., 2024). Prenatal acoustic environment was considered as monolingual (i.e., no months of exposure) when mothers reported speaking only one language during their last trimester of pregnancy, or as bilingual (i.e., three months of exposure) when they reported using two different languages during that period. Postnatal linguistic exposure was evaluated for their first year of life through the Language Exposure Assessment Tool (LEAT; DeAnda et al., 2016), which provided the total number of months of bilingual exposure. Prenatal and postnatal exposure were analyzed within a unique continuous variable counting the number of months of bilingual exposure along the studied perinatal period (i.e., from the third trimester of pregnancy up to one year post-birth; Austin, 2014; Chiera et al., 2020; Marriott et al., 2019).

Bilingual exposure in the sample was characterized by the combination of Spanish and other language, being for most of the cases the Spanish-Catalan combination (89.3%). Other languages heard by infants were Bulgarian, English, Galego, Guarani, Moroccan Arabic and Portuguese (see more details in Table 1). Languages composing monolingual environments in the sample were either Spanish (88.2%) or Catalan (11.8%).

**Table 1.** Languages heard by infants from the third trimester of pregnancy to 12 months of age.

| Languages | N | % |
| --- | --- | --- |
| Spanish | 15 | 20.5 |
| Catalan | 2 | 2.7 |
| Spanish-Catalan | 49 | 67.1 |
| Spanish-Other | 7 | 9.7 |
| Bulgarian | 1 | 1.4 |
| English | 1 | 1.4 |
| Galego | 1 | 1.4 |
| Guarani | 2 | 2.7 |
| Moroccan Arabic | 1 | 1.4 |
| Portugal | 1 | 1.4 |

### 2.3 Stimulus

Neural responses were obtained to a 250 ms two-vowel /oa/ speech stimulus with a rising pitch ending, previously designed in our laboratory (for a detailed description, see Arenillas-Alcón et al., 2021). Three different sections can be differentiated in the stimulus according to its F_0_ and F_1_: the /o/ section (from 10 to 80 ms; F_0_ = 113 Hz; F_1_ = 452 Hz), the /a/ steady section (from 90 to 160 ms; F_0_ = 113 Hz; F_1_ = 678 Hz) and the /a/ rising section (from 160 to 250 ms; F_0_ = 113-154 Hz; F_1_ = 678 Hz). Both steady sections of the stimulus (/o/ and /a/ steady sections) were considered together for the analysis of the F_0_ encoding, giving rise to the stimulus steady section (from 10 to 160 ms; F_0_ = 113 Hz).

The specific vowels’ F_1_ were chosen as those belong to the prototypical phonetic repertoire in both Spanish and Catalan languages (Alarcos Llorach, 1965; Martí i Roca, 1986). The stimulus was presented at a rate of 3.39 Hz and an intensity of 60 dB SPL to the right ear. It was delivered in alternating polarities through an earphone connected to a Flexicoupler® disposable adaptor (Natus Medical Incorporated, San Carlos, CA).

### 2.4 Procedure and data acquisition

Neonatal FFR responses were recorded while the babies were sleeping in their crib at the hospital room. FFR sessions at six and twelve months of age were conducted at a hospital dispensary while ensuring the infant remained either asleep or as calm as possible, aiming to guarantee the highest quality of data. The recording sessions had a total mean duration of around 30 minutes, including 5 minutes of preparation time, 20 minutes of recording (4 /oa/ blocks × 1000 sweeps × 295 ms stimulus-onset asynchrony), and up to 5 minutes of additional time for the rejected sweeps.

The speech stimulus was presented using a SmartEP platform connected to a Duet amplifier, which includes the cABR and Advanced Hearing Research modules (Intelligent Hearing Systems, Miami, Fl, USA). Neural responses were recorded using three disposable Ag/AgCl electrodes placed in a vertical montage (active electrode located at Fpz, ground at the forehead and reference at the right mastoid), keeping impedances below 10 kΩ for all electrodes. The continuous FFR signal acquisition was completed at a sampling rate of 13333 Hz utilizing an online bandpass filter to eliminate frequencies outside the 30 to 1500 Hz range. EEG online data was epoched from -40.95 (pre-stimulus period) to 249.975 ms, automatically excluding any sweep with voltage values exceeding ± 30 µV.

### 2.5 Data processing

Acquired FFRs were bandpass filtered from 80 to 1500 Hz. To emphasize the FFR components associated to the speech stimulus envelope (FFR_ENV_) and to diminish putative cochlear microphonics, neural responses to alternating polarities were averaged [(Condensation + Rarefaction)/2]. To assess the neural encoding of the stimulus’ F_1_, and minimizing the envelope related neural activity (Aiken & Picton, 2008; Krizman & Kraus, 2019), the FFR temporal fine structure (FFR_TFS_) was obtained by subtracting neural responses to the two opposite polarities [(Rarefaction–Condensation)/2].

FFR parameters were estimated using custom scripts from Matlab R2019b (The MathWorks Inc., 2019), previously developed in our laboratory and used in former similar studies (Arenillas-Alcón et al., 2021; Ribas-Prats et al., 2019). A detailed description can be found below for the three FFR parameters separately extracted and tested for the different stimulus features of interest.

#### Spectral amplitude

Spectral amplitude (in nV) was obtained as an indicator of the neural-phase locking magnitude at the frequency of interest (F_0_, 113 Hz; /o/ F_1_, 452 Hz; /a/ F_1_, 678 Hz) (Arenillas-Alcón et al., 2021; Ribas-Prats et al., 2019; White-Schwoch et al., 2015b). It was calculated by applying the Fast Fourier Transform (FFT; Cooley & Tukey, 1965) to the neural response. Spectral amplitude was defined as the mean amplitude within a ±5 Hz window centered at the frequency peak of interest. Spectral amplitude at F_0_ was obtained from the FFR_ENV_ corresponding to the stimulus steady section (10 to 160 ms) to quantify voice pitch encoding of the speech-sound stimulus. Spectral amplitudes at the vowels’ F_1_ frequencies were retrieved separately from the FFR_TFS_ corresponding to the /o/ section (10 to 80 ms) and the /a/ steady section (90 to 160 ms).

#### Pitch error

Pitch error (in Hz) was extracted from the FFR_ENV_ as a measure of pitch encoding accuracy for the F_0_ contour along the two /a/ sections of the stimulus (i.e., /a/ steady section and /a/ rising section). It was computed as an average of the absolute Euclidian distance between the stimulus and response F_0_ from each bin separately for the two sections mentioned.

### 2.6 Statistical analysis

Statistical analyses were performed using Jamovi 2.4.11 (The Jamovi Project, 2024). To explore the effects of perinatal bilingual environment exposure and sex on the developmental trajectory of neural encoding of speech, linear mixed effects models were constructed separately for each FFR parameter according to our hypothesis: spectral amplitude (at 113 Hz, 452 Hz and 678 Hz) and pitch error (during the /a/ steady and /a/ rising). Normality was assessed with Kolmogorov-Smirnov test and a natural logarithm (ln) transformation was applied to spectral amplitude dependent variables, as those did not meet such assumption. Five different models were created to explore the trajectory of neural encoding of speech sound characteristics, one per each dependent variable, as follows:

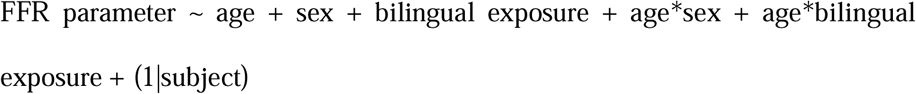

Individual tested models predicted each FFR value as a function of age (birth, six and twelve months) and sex (male, female) and bilingual perinatal exposure (as a covariate, ranging from 0 to 15 months of exposure). The models also included the interaction effect of age per sex, and age per bilingual exposure. A by-subject random intercept was included in the models to account for infants’ repeated measures.

A trend analysis was performed within each separated model by coding the age variable with polynomial contrasts, which describe possible trends in the means (i.e., shape of the age-dependent trend). The polynomial regression was thus applied to depict the relationship between the FFR parameters and age to find the best way to draw a line through the data points. After a significant result for the omnibus test of fixed effects, post-hoc ANOVA or multiple comparison with Bonferroni adjustments were conducted to further inspect the given result. Only Bonferroni-corrected p-values are reported in the results section. Normality of residuals was met for each independent model, which was checked using Kolmogorov-Smirnov test.

## 3. Results

Clear FFRs were elicited across each developmental stage (i.e., at birth, six months and twelve months) in both female and male infants. These neural responses are illustrated in Fig.1, as FFR waveforms and corresponding spectral representations. As it can be observed in the figure, spectral peaks are present at F_0_ across ages, and for the stimulus F_1_ emerging from 6 months onwards.

**Fig. 1.**
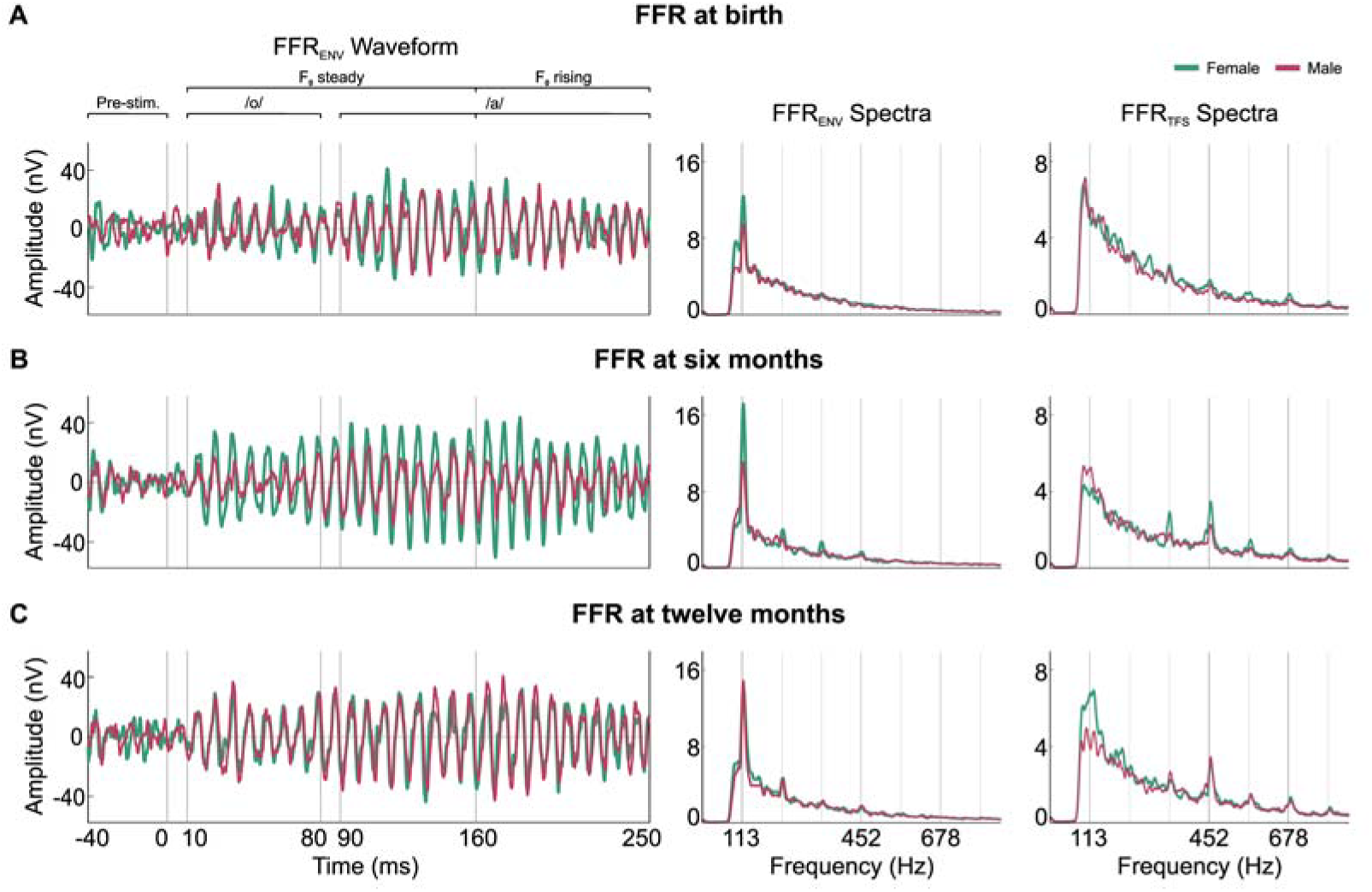
FFRs from 38 female (in green) and 35 male infants (in red) recorded at each developmental stage: (**A**) at birth, (**B**) at six months and (**C**) at twelve months. The left column displays grand-averaged FFR_ENV_ waveforms in the time domain for each age group. The middle and right columns illustrate the amplitude spectra of FFR_ENV_ and FFR_TFS_, respectively, both extracted from the steady section of the stimulus (10–160 ms). Remarkably, a clear sex difference in F_0_ encoding emerges at six months of age, with females displaying larger amplitudes than males.

### 3.1 Voice pitch encoding

#### Spectral amplitude at stimulus F_0_

The model fit was statistically significant (*X*^2^(9) = 37.544; R^2^ = .275; *p* < .001). The regression results for the model indicated a main effect of sex (β = -.242; t(70.6) = -2.312; *p* = .024), with general higher spectral amplitudes shown by female infants (*M* = .013 ± .008) in comparison to their male peers (*M* = .01 ± .006; see Fig. 2A). The interaction effect between age and sex was found to be a significant predictor (*F*_(2,140.3)_ = 3.15; *p* = .046), with higher values for female in comparison to male infants at the age of six months (t(199) = 3.36; *p* = .014). Moreover, the interaction between the quadratic effects of age and sex was found as a significant predictor of the spectral amplitude at 113 Hz (β = .347; t(140.2) = 2.43; *p* = .016). Simple slopes analysis revealed that the quadratic trajectory of spectral amplitude as a function of age was only present for female infants (β = -.237; t(156) = -2.01; *p* = .047). The interaction effect between age and bilingual exposure was also significant (*F*_(2,173.6)_ = 8.19; *p* < .001; see Fig. 2A). Simple slope analysis revealed a negative effect of bilingual exposure on spectral amplitudes at six months (β = -.051; t(209.4) = -2.42; *p* = .017), along with a positive effect at twelve months (β = .043; t(209.9) = 3.27; *p* = .001). The quadratic effect of age per bilingual exposure interaction was also found to be significant (β = .062; t(161.1) = 2.33; *p* = .021). Simple slopes analysis revealed that only for monolingual infants (i.e., none-exposed to bilingualism) a significant quadratic effect of age was predicted on spectral amplitude at 113 Hz (β = -.340; t(142.1) = -3.06; *p* = .003).

**Fig. 2.**
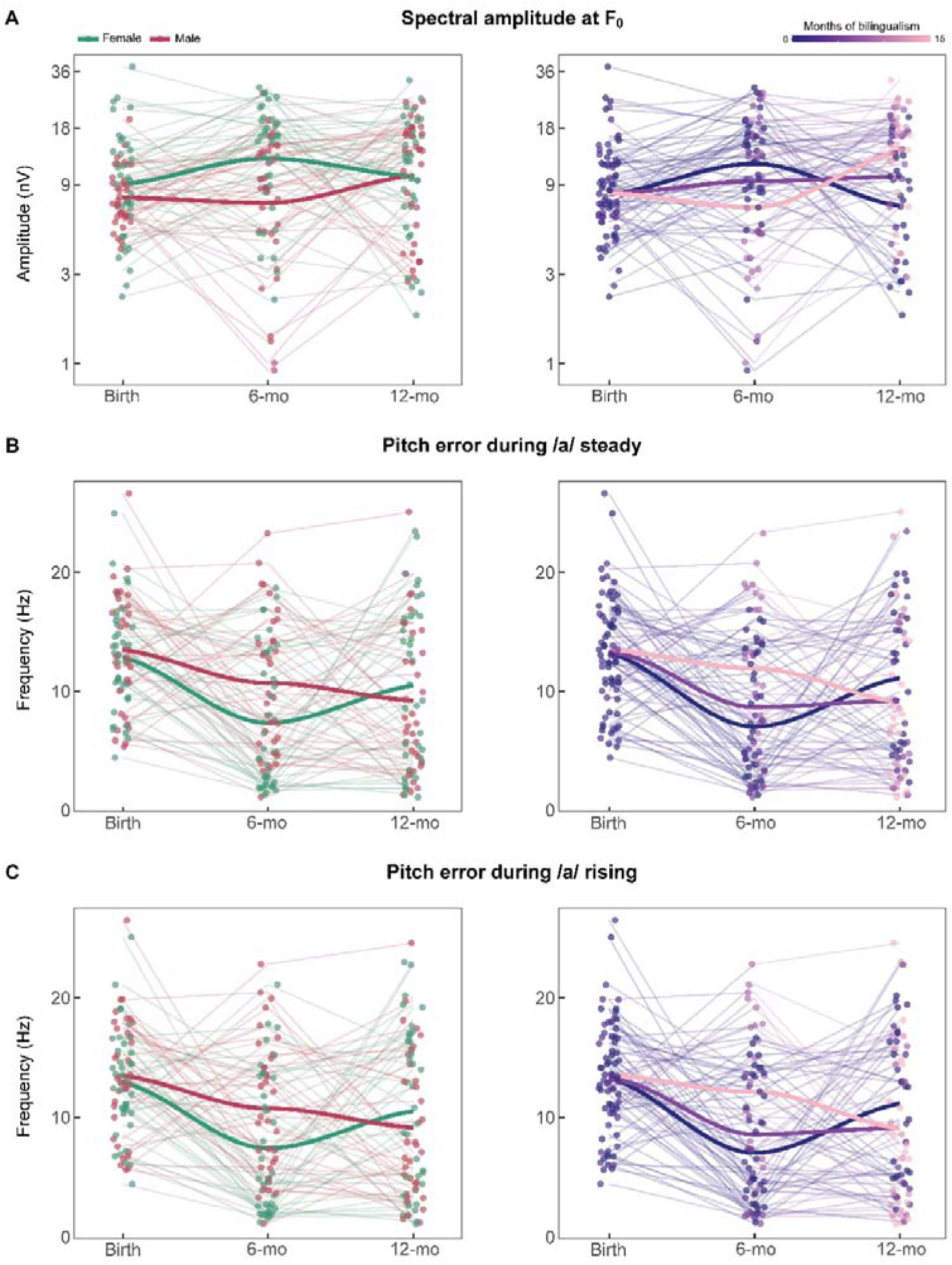
Predicted developmental trajectories of (**A**) spectral amplitudes at F_0_ and (**B**) pitch error during both /a/ steady and (**C**) rising sections of the stimulus across the three developmental stages: at birth, six months (6-mo) and twelve months (12-mo). Solid lines illustrate the predicted trajectories based on sex (left panel: female in green, male in red) and bilingual exposure (right panel: no exposure in dark blue, median exposure in purple, maximum exposure in pink). (**A**) Each data point represents the log-transformed spectral amplitudes of the longitudinally tested infants to account for skewness. For interpretability, the y-axis displays the corresponding real spectral amplitude values. Data points displayed in (**B**) and (**C**) represent the non-transformed pitch error values corresponding to each tested infant.

#### Pitch error during the /a/ steady section

The model fit was statistically significant (*X*^2^(9) = 38.618; R^2^ = .232; *p* < .001). The regression results for the model indicated a main effect of age (*F*_(2,176.9)_ = 3.58; *p* = .030), with higher values at birth (*M* = 13.25 ± 1.43) in comparison to six months of age (*M* = 9.19 ± .66; t(194.1) = 2.64; *p* = .027) and depicting a significant quadratic trajectory (β = 2.09; t(162.9) = 2.55; *p* = .012). The interaction between the quadratic effect of age and sex was found as a significant predictor of pitch error during the /a/ steady section (β = -2.826; t(140.5) = -2.25; *p* = .026; see Fig. 2B), with simple slopes analysis revealing a quadratic trajectory only significantly present for female infants (β = 3.504; t(156.8) = 3.39; *p* < .001). The interaction effect between age and bilingual exposure was also found to be a significant predictor (*F*_(2,175.1)_ = 4.31; *p* = .015; see Fig. 2B). Post-hoc estimates revealed higher pitch error values as a function of bilingual exposure at six months of age (β = .470; t(209.7) = 2.62; *p* = .009).

#### Pitch error during the /a/ rising section

The model fit was statistically significant (*X*^2^(9) = 39.642; R^2^ = .241; *p* < .001). The regression results for the model indicated a main effect of age (*F*_(2,176.9)_ = 3.51; *p* = .032), with higher values at birth (*M* = 13.28 ± 1.42) in comparison to six months of age (*M* = 9.25 ± .66; t(194) = 2.62; *p* = .028) and depicting a quadratic effect of age as a significant predictor of pitch error during the /a/ rising section (β = 2.04; t(162.7) = 2.50; *p* = .013). The interaction between age and sex was found as a significant predictor (*F*_(2,140.6)_ = 3.14; *p* = .046). Post-hoc comparisons revealed higher values for female infants at six months in comparison to male neonates (t(209.8) = 3.35; *p* = .014). The interaction between the quadratic effects of age and sex was also found as a significant predictor (β = -2.871; t(140.5) = -2.30; *p* = .023; see Fig. 2C). Simple slopes analysis revealed that the quadratic trajectory of pitch error as a function of age was only present for female infants (β = 3.474; t(156.7) = 3.38; *p* < .001). The interaction effect between age and bilingual exposure was also found to be a significant predictor (*F*_(2,175)_ = 4.72; *p* = .010; see Fig. 2C). Post-hoc estimates revealed higher pitch error values as a function of bilingual exposure at six months of age (β = .491; t(209.7) = 2.75; *p* = .006).

### 3.2 Formant structure encoding

#### Spectral amplitude at /o/ vowel F_1_

Model fit was statistically significant (*X*^2^(9) = 30.512; R^2^ = .211; *p* < .001). The regression results for the model indicated a main effect of age (*F*_(2,176.6)_ = 4.77; *p* = .01), with lower values at birth (*M* = .003 ± .003) in comparison to both six months (*M* = .004 ± .004; t(193.8) = -2.93; *p* = .011) and twelve months of age (*M* = .004 ± .004; t(208.2) = -2.95; *p* = .010). Both a linear (β = .526; t(208.2) = 2.96; *p* = .003) and a quadratic (β = -.276; t(162.2) = -2.15; *p* = .033) effect of age resulted as significant predictors of spectral amplitude at 452 Hz. The interaction between age and sex was found to a be significant predictor (*F*_(2,140.2)_ = 6.15; *p* = .003; see Fig. 3A), with six-months old females showing higher spectral amplitudes in comparison to both male (t(210) = -3.27; *p* = .019) and female neonates (t(184) = -3.83; *p* = .003). Significantly higher spectral amplitudes were also depicted for twelve-months male infants in comparison to female neonates (t(204) = -3.23; *p* = .022). The interaction of age as following a quadratic trajectory by sex was found as a significant predictor (β = .684; t(140.0) = 3.48; *p* < .001), with post-hoc results showing a significant quadratic effect of age only for female participants (β = -.618; t(156) = -3.81; *p* < .001).

**Fig. 3.**
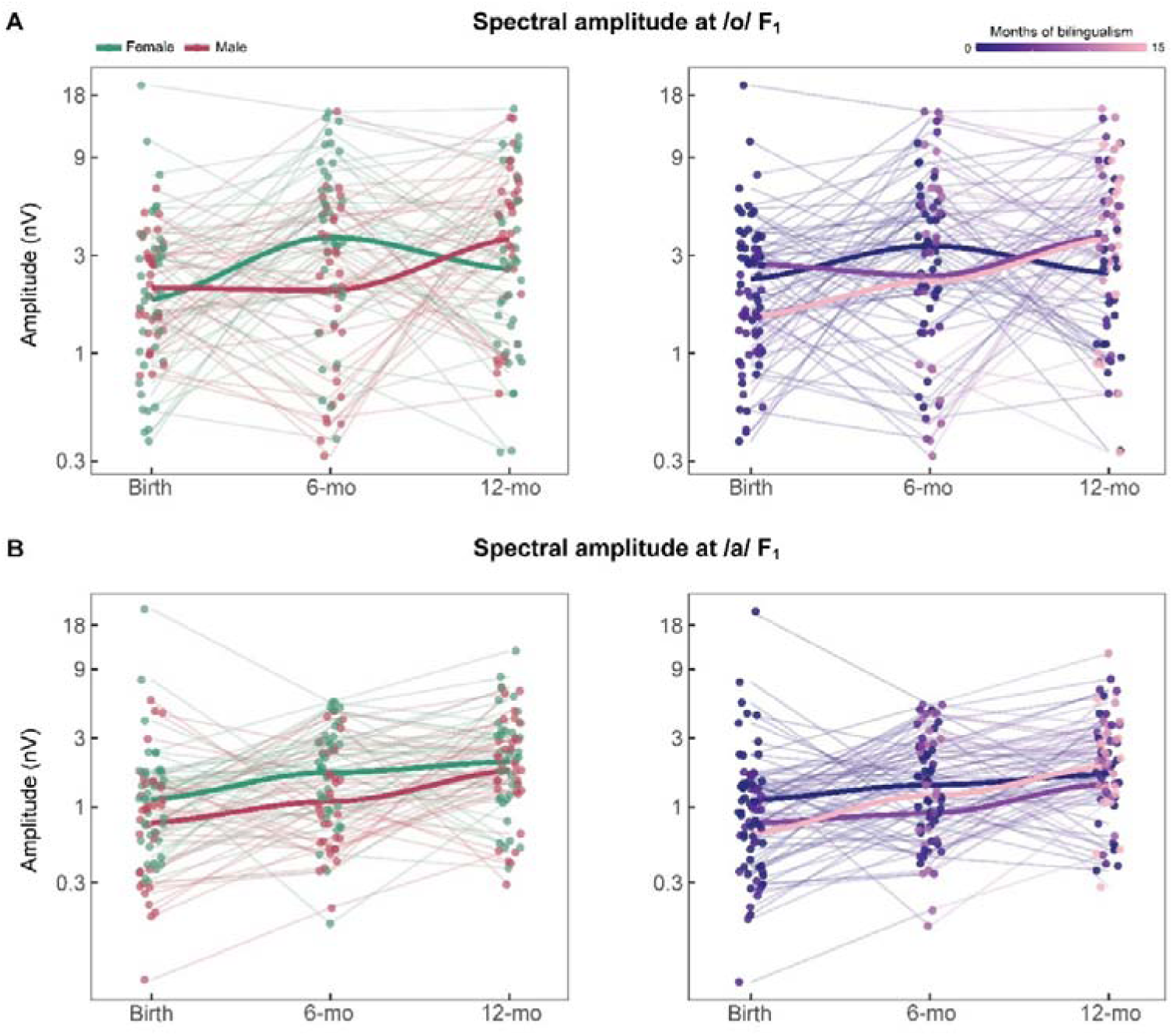
Predicted developmental trajectories of neural encoding for (**A**) /o/ F_1_ and (**B**) /a/ F_1_ across the three developmental stages: at birth, six months (6-mo) and twelve months (12-mo). Each point represents the log-transformed amplitudes for each longitudinally tested infant to account for skewness. For interpretability, the y-axis displays the corresponding real spectral amplitude values. The solid lines represent the predicted trajectories according to sex (left panel: female in green, male in red) and bilingual exposure (right panel: no exposure in dark blue, median exposure in purple, maximum exposure in pink).

The interaction between age and bilingual exposure was also significant (*F*_(2,174.6)_ = 3.41; *p* = .035; see Fig. 3A), although post-hoc estimates results did not reach statistical significance at any developmental stage. The linear effect of age per bilingual exposure interaction was also found as a significant predictor (β = .115; t(199.3) = 2.26; *p* = .025), indicating that the linear trajectory of age on spectral amplitude at 452 Hz depends on the level of bilingual exposure. Simple slopes analysis revealed a predicted linear effect of age specifically for infants with medium level of bilingual exposure (7.5 months of exposure; β = .920; t(207.8) = 2.81; *p* = .005) and for those with maximum bilingual exposure (15 months of exposure; β = 1.780; t(204.7) = 2.56; *p* = .011).

#### Spectral amplitude at /a/ vowel F_1_

Model fit was statistically significant (*X*^2^(9) = 47.423; R^2^ = .323; *p* < .001). The regression results indicated a main effect of age (*F*_(2,176.7)_ = 10.75; *p* < .001), with lower values at birth (*M* = .0015 ± .003) in comparison to both six months (*M* = .0018 ± .001; t(193.1) = -3.53; *p* = .002) and twelve months of age (*M* = .0025 ± .002; t(208.4) = -4.61; *p* < .001). Moreover, a linear effect of age was depicted as a significant predictor of spectral amplitude at 678 Hz (β = .769; t(208.4) = 4.61; *p* < .001). A main effect of sex was also discovered (β = -.309; t(70.9) = -2.28; *p* = .026; see Fig. 3B), with general higher values for female infants (*M* = .0022 ± .003) than their male peers (*M* = .0016 ± .001). The linear effect of age per bilingual exposure interaction was also found as a significant predictor (β = .111; t(196.2) = 2.36; *p* = .019; see Fig. 3B), specifically indicating that the linear trajectory of age on spectral amplitude at 678 Hz depends on the level of bilingual exposure. Simple slopes analysis revealed a predicted positive linear effect of age for all three contrasted levels of bilingual exposure: none exposed infants (β = .317; t(159.6) = 2.40; *p* = .018), infants exposed during 7.5 months (β = 1.151; t(206.9) = 3.75; *p* < .001), and infants exposed during 15 months to bilingual exposure (β = 1.986; t(202.8) = 3.07; *p* = .002).

## 4. Discussion

The present study was set to investigate the distinct trajectories in the neural encoding of speech sound features across the first year of infant development, with a focus on the influences of sex and perinatal bilingual exposure. To that aim, we inspected FFR neural responses to a two-vowel /oa/ syllable at birth, six months, and twelve months of age. We then modeled the neural encoding of both pitch and formant structure over time, analyzing variations according to bilingualism and sex. Our findings provide new insights into how sex and early bilingual exposure shape the neural encoding of speech sounds in infancy, with potential implications for understanding the individual variability in early language acquisition.

Our results reveal a significant maturation of neural encoding for both voice pitch and formant structure of speech during the first six months of life, followed by a stabilization during the second half of the year. Notably, different developmental trajectories were observed for each of the different speech features of interest. While the encoding of both steady and rising F_0_ contours followed a quadratic trajectory, the encoding of both vowels’ F_1_ exhibited a linear progression. Interestingly, the encoding of the /o/ F_1_ frequency fit both linear and quadratic models, suggesting a more complex developmental pattern. These divergent trajectories may reflect differences related to the development of low- and high-frequency acoustic information processing, paralleling the spectrally ascendant developmental pattern of the auditory system (Graven & Browne, 2008). Accordingly, neural attunement to low frequency acoustic content initiates earlier, being the period from 25 gestational weeks to six postnatal months the most critical for the neurosensory development of the auditory system. This aligns with the earlier availability of low-frequency acoustic signals accessible to the fetus during the prenatal period (Hepper & Shahidullah, 1994; Voegtline et al., 2013) and is supported by previous research demonstrating robust pitch encoding already at birth (Arenillas-Alcón et al., 2021; Jeng et al., 2011). Intriguingly, although our findings did not reveal significant age-related changes in spectral amplitudes at F_0_, we did identify marked improvements in the neural tracking of both steady and rising pitch contours within the first six months. These results may help clarify inconsistencies in previous research on infant age-related changes in pitch encoding, which may stem from subtle variations in the features examined (Anderson et al., 2015; Jeng et al., 2010; Puertollano et al., 2024; Ribas-Prats et al., 2023b; Van Dyke et al., 2017).

On the contrary, the neural encoding of F_1_ components has been documented to experience significant enhancement already in the first postnatal month (Ribas-Prats et al., 2023b), suggesting the rapid neural adaptation to novel frequencies that become significantly more available after birth. Our results corroborate previous research demonstrating further refinement of F_1_ neural encoding during the first six months of life (Ribas-Prats et al., 2023b) and its stabilized development up to the age of twelve months (Puertollano et al., 2024), but further extend these findings by revealing a linear maturational pattern throughout the first year of life. This underscores the early development of neural mechanisms crucial for establishing a native language’s sound map by six months, thereby facilitating native phoneme identification and discrimination (Kuhl, 1991, 2004, 2010; Kuhl et al., 1992).

A complementary explanation for this early maturation pattern lies in the constraints imposed by neuronal development, as shown by the evolution of electrophysiological brain activity in early infancy. Fetal and neonatal brain activity is primarily characterized by slow-wave oscillations that match the slow prosodic modulations found in speech. By around six months of age, faster neuronal oscillations that can phase-lock to phoneme-rate amplitude modulations emerge (Anderson & Perone, 2018; Le Van Quyen et al., 2006; Xu et al., 2011; for a review see Menn et al., 2023), enabling infants to encode higher-frequency speech components and build a repertoire of native phonemes (Kuhl, 2004). Neural encoding of phonetic features becomes robust by seven months of age and shows no further enhancement up to the age of eleven months (Di Liberto et al., 2023). Similarly, our findings highlight the first six months as a critical period, when the neural system increasingly attunes to the spectro-temporal features of the acoustic environment along the auditory pathway, setting the stage for subsequent phonetic learning (Werker & Hensch, 2015; Zhao et al., 2022).

There is a vast body of literature emphasizing the complex interplay between maturational and experiential influences on speech development (for a review see Werker & Hensch, 2015). Evidence for a sensitive period for phonetic attunement occurring during the first twelve months of life comes from diverse biological and environmental contexts. While studies on infants born prematurely emphasize the relevance of maturational processes in speech perception development (Peña et al., 2010, 2012), research into acoustic deprivation due to infant deafness underscores the critical role of sensory input in driving acoustic-related neural plasticity during the first year of life (Faulkner & Pisoni, 2013; Kral & Sharma, 2012). A particular acoustic environment is that related to bilingual experience, which has been previously proposed to modulate the onset and duration of the sensitive period for phonetic learning (Best et al., 2016; Garcia-Sierra et al., 2011; Werker et al., 2009; Werker & Hensch, 2015). This extended sensitive period likely offers bilingual infants additional time to attune to the acoustic and phonological features of both languages. Our findings support this proposal by underscoring the moderating influence of bilingualism on the development of neural mechanisms involved in pitch and formant structure encoding, both of which are essential for speech and language acquisition.

We observed a negative impact of bilingual exposure on voice pitch encoding at six months, as reflected by lower spectral amplitudes at F_0_ and higher pitch error values. This trend reversed by twelve months, with bilingual exposure predicting higher spectral amplitudes at F_0_. This U-shaped pattern could reflect an initial struggle to attune to speech sounds, related to a more complex acoustic environment. Notably, immature neural responses to phonemic contrasts during the early months of life do not necessarily predict poorer language outcomes. Instead, they seem to facilitate latter attunement, leading to faster maturation rates and stronger responses later in development that are associated to enhanced language outcomes (García-Sierra et al., 2021; Schaadt et al., 2015, 2023; Werwach et al., 2022). In contrast, monolingual infants in our sample followed an inverted U-shaped trajectory for F_0_ encoding, with spectral amplitudes increasing from birth to six months and then declining between six and twelve months. These opposite trajectories in F_0_ encoding depending on bilingual experience parallel those depicted for phonetic contrast discrimination during the second half of the first infant year (Byers-Heinlein & Fennell, 2014). F_0_ variability may serve as a salient perceptual attribute that facilitates language separation for bilingual infants (Liu & Kager, 2017).

Bilingual experience further affected the maturation of formant structure encoding, with distinct effects on each vowel’s F_1_ frequencies. Specifically, infants with over seven months of total bilingual exposure showed linear growth in spectral amplitudes for /o/ F_1_ across the first year, whereas this linear effect of age was not significant for monolingual infants. In contrast, neural encoding of /a/ F_1_ followed a linear trajectory regardless of the amount of bilingual exposure. Our findings suggest a stronger influence of bilingual exposure in lower versus higher frequencies during early infant ages that may be related to the previously mentioned spectrally ascendant pattern of development (Graven & Browne, 2008). However, although describing distinct developmental trajectories for the encoding of pitch and formant structure, our results demonstrate that bilingual exposure positively impacts the neural encoding of both speech features at the age of twelve months. We thus provide evidence for previous literature hypothesizing a heightened acoustic sensitivity in bilingual infants manifesting near the end of the perceptual reorganization process (Liu & Kager, 2015, 2017), by specifically signaling distinct neural attunement along the auditory pathway associated to early bilingual exposure. Additionally, our results complement previous research on the effects of prenatal bilingual exposure in neural encoding of speech sounds (Gorina-Careta et al., 2024) by extending the scope to postnatal development throughout the first year of life.

Our findings further underscore a significant influence of sex on the neural speech encoding in infants. Female infants exhibited higher overall spectral amplitudes at the stimulus F_0_, indicating superior voice pitch neural phase-locking throughout the first year compared to males. Additionally, females followed a distinct quadratic developmental trajectory for pitch encoding, surpassing male spectral amplitudes at F_0_ by six months and then declining closer to male values by twelve months. A similar quadratic pattern was observed for females in the encoding of /o/ F_1_, marked by significant improvements during the first six months of life. Female infants also displayed overall higher amplitudes at /a/ F_1_ across the first year. These results extend those of Krizman et al. (2019, 2020) by revealing sex differences in auditory processing emerging in infancy, well before the adolescent differences they documented.

These sex-specific developmental patterns highlight early differences in neural encoding of speech-acoustic features between male and female infants during the first year of life. Our results align with previous research reporting sex differences in early speech encoding and acquisition, including disparities in the maturation of involved brain areas (Alexopoulos et al., 2022), infant phonological discrimination (Friederici et al., 2008), the onset and rate of word production (Dailey & Bergelson, 2023), and the frequency of speech-like vocalizations before first words (Sung et al., 2013). A growing body of research links infants’ hormone levels to these early developmental differences (Lust et al., 2010; Lutchmaya et al., 2001; Wermke et al., 2014). For example, sex hormone levels have been linked to infant phonological discrimination (Friederici et al., 2008), articulatory skills (Quast et al., 2016), and later language abilities in childhood (Hollier et al., 2013; Schaadt et al., 2015).

Another factor potentially contributing to these differences is the related to caregivers’ speech input. Studies suggest that caregivers use higher pitch and a greater pitch range when speaking to female infants compared to males, with this difference growing over time (Kitamura et al., 2001; Kitamura & Burnham, 2003). The duration of speech input also shows sex-specific patterns, with speech directed to male infants decreasing during the first year of life, while increasing for female infants (Sung et al., 2013). Furthermore, caregivers tend to talk more to infants who have started talking (Dailey & Bergelson, 2023), which could reinforce sex differences in speech abilities given that females often begin speaking earlier than males (Bornstein et al., 2004; Eriksson et al., 2012; Fenson et al., 1994). These biological and social factors likely interact, shaping the sex-related differences in the developmental trajectory of neural speech encoding observed in our results.

Although there is a large research community exploring speech perception in infants, using both behavioral and neurophysiological measures (Byers-Heinlein et al., 2010; Hervé et al., 2022; Kuhl et al., 2003, 2006; Kujala et al., 2023), the use of the FFR provides a direct neural measurement of the auditory hierarchy that reflect the attunement to the spectral and temporal features of speech (Coffey et al., 2016; Gorina-Careta et al., 2021; Krizman & Kraus, 2019; Skoe & Kraus, 2010). By utilizing the FFR, our results highlight the pivotal importance of the first six months in neural development for speech encoding. Thereby, future studies should incorporate more frequent time-point measurements to gain a comprehensive understanding of this early period. We observed distinct developmental trajectories in the neural encoding of speech features, influenced by both sex and perinatal bilingual exposure. Consistently incorporating these factors into research on early speech and language development is crucial for capturing an accurate depiction of these early processes, enhancing replicability and deepening our understanding of infant development.

## 5. Conclusion

The present study provides novel insights into the distinct developmental trajectories of neural speech encoding during the first year of life, influenced by both sex and perinatal bilingual exposure. By examining longitudinally FFR neural responses to a two-vowel stimulus, we revealed a significant maturation of pitch and formant structure encoding throughout the first six months of life, without further maturation up to the age of twelve months. These findings emphasize the first six months as a critical period for neural adaptation to the acoustic environment, contributing to early phonetic learning and language acquisition. The moderating effects of bilingual exposure on voice pitch and formant encoding, as well as the sex-specific differences observed, underscore the complexity and individual variability related to early auditory and speech perception development. These results not only extend previous research but also contribute to a deeper understanding of the neural foundations of early language acquisition. Future studies incorporating more frequent developmental assessments could help refine our understanding of this sensitive period.

## Data Availability Statement

The data that support the findings of this study are available from the corresponding author Carles Escera upon request.

